# Timing of mortality during development alters the evolution of aging

**DOI:** 10.1101/2023.01.24.525295

**Authors:** Peter Lenart, Sacha Psalmon, Benjamin D. Towbin

**Affiliations:** University of Bern, Institute of Cell Biology, Bern, Switzerland; Polytech Nice Sophia, Côte d’Azur University, Nice, France

**Keywords:** Aging, development, mortality rates, extrinsic mortality, negligible senescence

## Abstract

**Background:** For most animals, intrinsic senescence-induced mortality increases with age, while deaths from extrinsic threats, such as predation or accidents, decline during development as individuals grow and mature. Age-dependent modulation of extrinsic mortality is known to influence the evolution of aging, yet how the timing of mortality shapes evolutionary forces remains poorly understood.

**Results:** To address this, we developed two complementary mathematical models that integrate survival benefits arising during development with the progressive increase in mortality associated with senescence. Agent-based simulations and deterministic analyses revealed a strong and consistent influence of the timing of developmental survival benefits on their evolutionary impact: early-life survival benefits reduced the selection for slower aging, while late-acting benefits enhanced it. This difference arises because early-life benefits more strongly accelerate population growth than late-life benefits, diminishing the relative evolutionary advantage of increased longevity.

**Conclusions:** Our results underscore the importance of mortality timing in the evolution of aging and provide a theoretical framework for connecting developmental trajectories to aging dynamics.

## Background

Differences in developmental programs produce a wide range of age-specific mortality patterns among species (1). In many animals, the risk of death from extrinsic causes, such as predation or accidents, declines during development as individuals grow and mature (2–5). Conversely, as animals age, their risk of death caused by intrinsic failure increases due to progressive physiological decline (6,7). The interaction of mortality modulation through development and through aging is central to understanding the evolution of aging rates (8,9).

Classic theory by George Williams (10) emphasized a role for extrinsic mortality in shaping the evolution of aging (11,12). Williams argued that high extrinsic mortality weakens the selection for slower aging because most individuals die before reaching old age (10). Building on this verbal argument, subsequent mathematical models showed that this effect only occurs under certain population density-dependences (8,11,13–17) or when extrinsic mortality is age-dependent (14,16,17). These models also showed that age-dependent modulation of mortality can, in principle, either strengthen or weaken the selection for slower aging, but the relation between the time-dependence of mortality and the evolutionary consequence on aging remains incompletely understood.

Age-dependence of extrinsic mortality was already implicitly considered by Williams, who distinguished between adult and juvenile mortality (10). Importantly, however, extrinsic mortality can also vary after reaching adulthood. In humans, for example, mortality from accidents sharply declines in young adults due to behavioral shifts and reduced risk-taking (18,19). Other adaptive traits, such as strengthened immunity, can similarly reduce extrinsic mortality with age. These age-dependent modulations of mortality are a key aspect of diverse developmental strategies and are intimately connected to the evolution of aging rates.

Notably, the distinction of extrinsic and intrinsic mortality is not always self-evident (20), as mortality from the same external threat is often shaped by both developmental processes and age-related decline. For example, the ability to evade predators can improve during development due to muscle growth, but deteriorate in old age due to aging-related sarcopenia. In this study, we define changes in extrinsic mortality as age-related modulation of mortality risks that occurs independent of physiological senescence (13). By focusing on the processes that modulate the mortality risk, rather than on the ultimate cause of death, this definition resolves the above-mentioned ambiguity and allows us to explore how developmental processes influence the evolution of aging rates. Under this definition, extrinsic mortality is indeed rarely age-independent: instead, it commonly declines during development as animals grow, mature, and adjust their behavior to environmental challenges (11,20–22).

To explore how developmental changes in survival shape the evolution of aging, we developed two complementary mathematical models, which we analyzed by agent-based simulations and by deterministic calculations. Both models showed that the developmental timing of survival benefits is critical for their evolutionary impact: while early survival benefits reduce the selective pressure for slower aging, late survival gains enhance it. This pattern arises because early and late survival benefits differentially affect population growth, thereby altering the strength of selection for slow aging. Our results underscore the evolutionary significance of time-resolved, developmentally driven changes in mortality and offer a powerful framework for understanding the co-evolution of development and aging.

## Results

### Early survival benefits decelerate, and late benefits accelerate the evolution of slower aging in a minimal model

To investigate the interactions between aging and developmentally occurring survival benefits, we created a parsimonious model, called the Development-Aging (DA) model. The model assumes that mortality due to aging increases exponentially with time according to the Gompertz law (*h*_*G*_ *(t) =a*e*^*bt*^) (23), and that deviations from the Gompertz law are due to developmentally occurring survival benefits that modulate this mortality. For this minimal model, we describe survival benefits *β* as a step function that decreases mortality by 20% from age *d* onwards, yielding an overall mortality

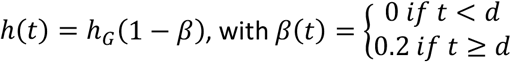

We call the parameter *b* the aging rate, i.e. the exponential rate of increase in the hazard rate if there were no developmentally occurring survival benefits.

To ask how the timing of developmental survival benefits affects the evolution of aging, we used agent-based simulations, where individual agents underwent repeated rounds of death, reproduction, and mutation. Simulations were carried out with a population of 10’000 individuals. The mortality rate of each individual was determined by the DA model, and each individual was given a 2% chance of mutation to alter the rate of aging (i.e., parameter *b*). Reproduction was sexual with constant fertility of one offspring per time step between ages 25 to 100 and zero fertility outside of this interval. Population density-dependence was implemented by filling freed spaces by random selection of offsprings created in the previous reproduction step. In this minimal model, we did not consider tradeoffs between aging and other life-history traits.

As expected, the aging rate rapidly declined during simulations (Fig. 1a) and eventually plateaued when the benefit of any additional slow-down of aging was counteracted by genetic drift (Supplemental Fig. 1). To ask how developmentally controlled survival benefits impacted the evolution of aging, we systematically varied the age *d* at which the reduction in mortality occurred. We then determined how this timing affected the rate of decline in the aging rate during simulations. For small *d*, i.e. an early onset, survival benefits decelerated the evolution of slower aging. However, with increasing *d* this effect became weaker and eventually turned to accelerate the evolution of slower aging (Fig. 1a, b). The effect size on the rate of evolution scaled with the magnitude of the survival benefit (parameter *β*), but the difference between early and late timing was qualitatively unaltered by this parameter (Fig. 1c, d).

**Figure 1.**
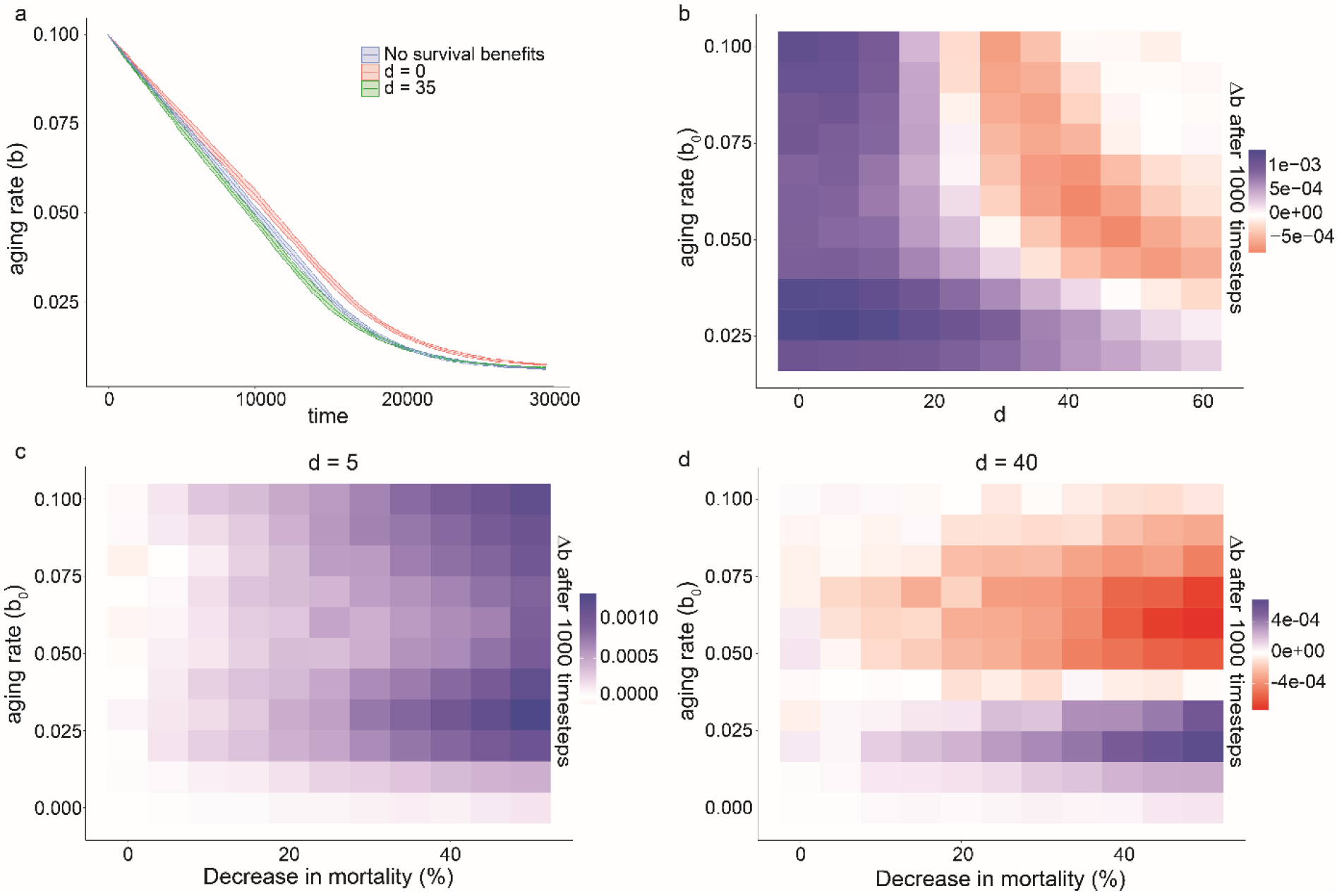
Early survival benefits decelerate, and late benefits accelerate the evolution of slower aging in the DA model. **a**. Evolution of the mean aging rate during agent-based simulations. Colors represent populations with timings of survival benefits early (red, *d* = 0) or late (green, *d* = 35) in life, compared to a population without developmentally controlled survival benefits (blue). The filled are on the plot shows 95% confidence interval of the mean. **b**. Difference in aging rate *Δb* within the first 1’000 simulation time steps between simulations starting at the same aging rate *b*_*0*_ with and without survival benefit (20%) occurring at age *d*. (*Δb* = (*b*_*0*_ – *b*_*1000, no benefit*_) - (*b*_*0*_ – *b*_*1000, benefit at age d*_) = *b*_*1000, benefit at age d*_ – *b*_*1000, no benefit*_)). Positive values (blue) indicate deceleration of evolution of slow aging by the survival benefit. Red indicates accelerated evolution. Each value represents the mean of 50 simulations of 10’000 individuals over 1’000 time points. **c**. same as **b**, but with a fixed value of *d* = 5 and different strengths of survival benefits. **d**. same as **c** but with *d* = 40

### Deterministic calculations confirm the conclusions from agent-based simulations

To validate the results from agent-based simulations, we computed the selective fitness as the intrinsic rate of population increase *r* by the Euler-Lotka equation (24) for a wide range of *b*_*0*_ and *d* of the DA model. These calculations closely matched results from agent-based simulations (Fig. 2), showing that the effect of developmental survival benefits on aging is not restricted to specific implementations used in the simulations, such as population density-dependence or population size.

**Figure 2.**
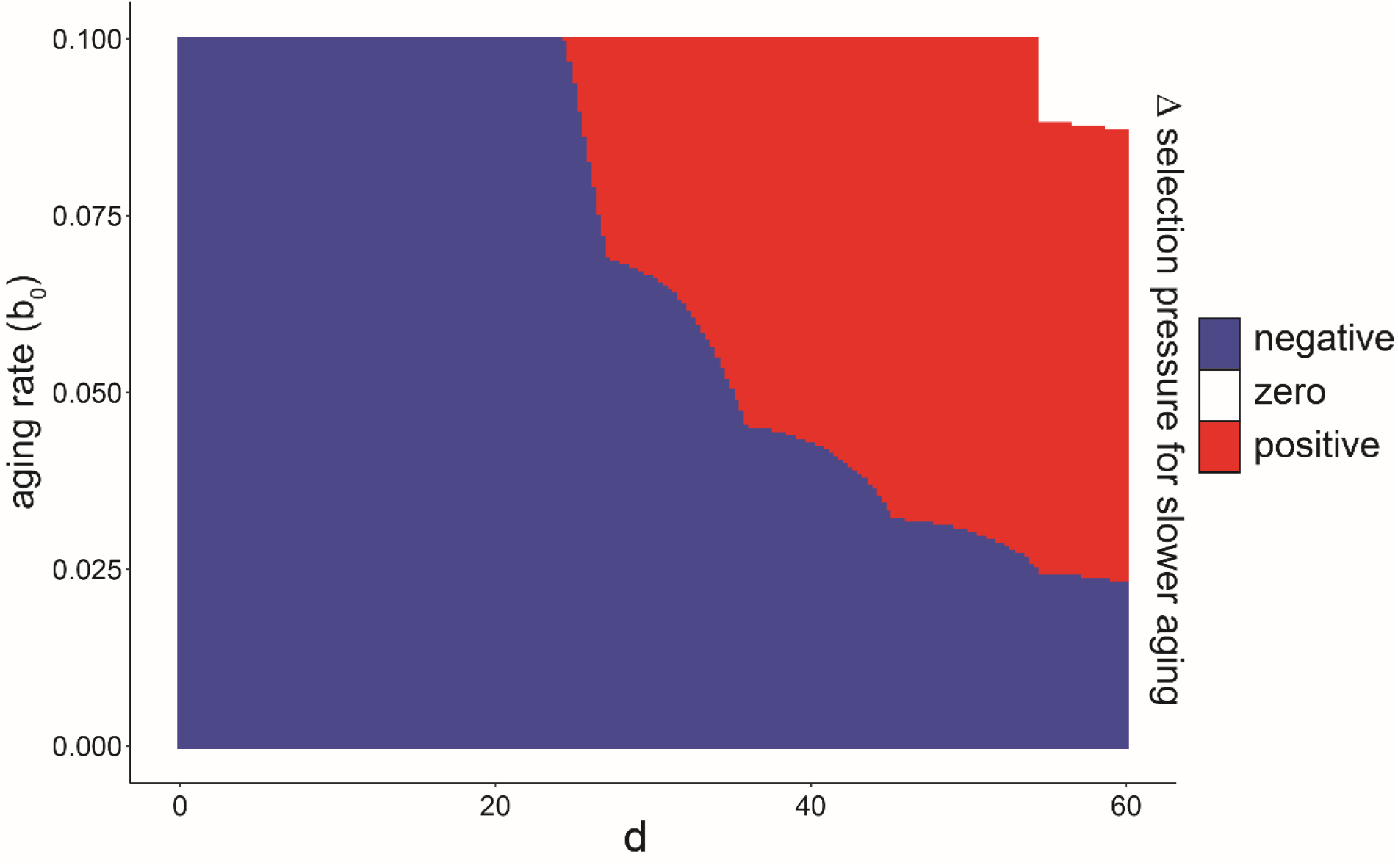
Deterministic calculations confirm the dependence of aging evolution on the developmental timing of survival benefits. Phase diagram distinguishing accelerating or decelerating effects of developmental survival benefits in the DA model calculated by the Euler-Lotka equation. The change in the intrinsic growth rate *Δr* after a 5% change in the aging rate *Δb* was calculated for life histories with and without survival benefits occurring at age *d* for different starting points of the aging rate *b*_*0*_. Blue area marks parameters in which the survival benefit decelerated the evolution of slower aging (*Δr* _*benefit at age d*_ > *Δr* _*no benefit*_). Red area: (*Δr* _*benefit at age d*_ < *Δr* _*no benefit*_). White area: 1^−13^ > (*Δr* _*benefit at age d*_ - *Δr* _*no benefit*_) > −1^−13^.

We conclude that, in the DA model, the timing of developmental survival benefits qualitatively changes their impact on the selection pressure for slower aging. The age *d* where selective pressure changed sign usually occurred around the age of first reproduction but also depended on the initial aging rate *b*_*0*_. This additional dependence suggests a more complex relationship between development and aging than a simple distinction between juvenile and adult mortality (25).

### An extended DA model describes deviations of human mortality from the Gompertz model by early and late survival benefits

To assess if these findings also apply to more complex mortality dynamics, we extended the DA model to capture realistic human mortality dynamics (human DA model, hDA). We chose human mortality dynamics for this analysis because of their complex shape, and for the unrivaled quality and availability of human mortality data. To capture the complex age-dependent mortality in humans, the hDA model uses the Gompertz-Makeham model (GMM), *h*_*GMM*_ *(t) = c + a*e*^*bt*^, and subtracts developmental survival benefits with either exponentially decelerating or logistic age dependence:

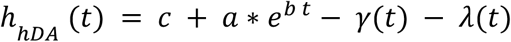

where 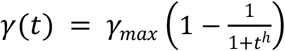 and 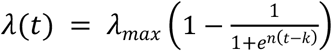

The terms *γ*_*max*_ and *λ*_*max*_ are scaling factors, corresponding to the maximal magnitude of the respective survival benefits. As for the DA model, we call parameter *b* of the Gompertz term the aging rate.

The hDA model provided good fits to human mortality data across juvenile and adult age classes for the population data from eight countries in the period from 1948 to 2020. Quality of fit was similar to other multi-parameter models of human mortality, such as Heligman’s model (26) (Fig. 3a-d). Specifically, the hDA model captured the rapid decline in mortality at a young age, and a plateau in the hazard rate during early maturity, as well as the exponential increase in mortality at ages above 30 years (Fig. 3a). As expected, the hDA model provided better fits than the GMM, which is not designed to capture all-age mortality, and the Siler model (27), which does not capture the mortality plateau in early adulthood (Fig. 3a).

**Figure 3.**
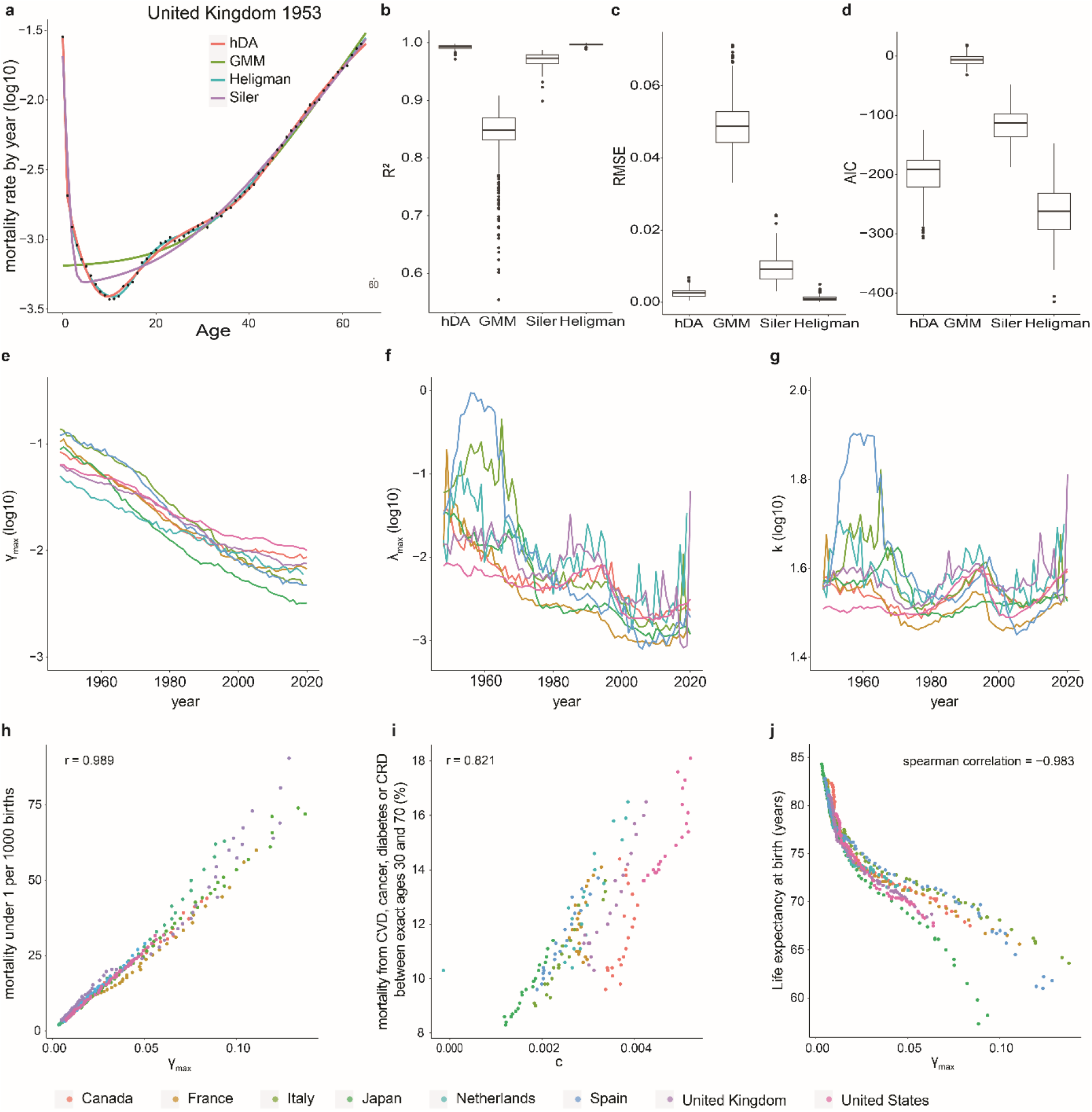
hDA model fits human mortality data. **a**. Fit of hDA (red), GMM (green) and Siler (purple), Heligman (blue) models to mortality data (black circles) from United Kindgom 1953 **b-d**. comparison of R^2^, RMSE and AIC calculated for DA, GMM, Siler, and Heligman models fitted to population data 1948-2022. Boxplot show 25^th^ - 75^th^ percentile. Line: median value. dots: outliers (>1.5 times the interquartile range away from the 25^th^ or the 75^th^ percentile). **e-g**. fitted values of *γ*_max_, *λ*_max_, and *k* of hDA model over time for indicated countries. **h**. correlation of *γ*_max_ and child mortality (29) for 1972-2020. **i**. correlation of *c* with mortality from cardiovascular disease (CVD), cancer, diabetes or chronic respiratory disease (CRD) (29) for 2000-2019. **j**. correlation of *γ*_max_ and life expectancy from 1948-2015. **e-j**. color represents countries, as indicated at the bottom of the figure.

The half-way points of *λ(t)*, determined by fits, ranged between 28 and 80, depending on year and country (mean = 36.56, sd = 7.22). The halfway point of *γ(t)* is inherently fixed to 1 year by the model. Thus, *γ(t)* represents early survival benefits, and *λ(t)* represents those occurring later in development. Although the hDA model is agnostic to the exact biological nature of the processes providing the benefit, these processes may involve developmental growth and developmental maturation for *γ(t)*, and adaptive immunity, behavioral changes, or learning for *λ(t)*.

Fitted parameters of the hDA model followed consistent trends across countries and time (Fig. 3e-g, Supplemental Fig. 2) that matched intuitive expectations. For example, *λ*_*max*_ and *γ*_*max*_ continuously declined between 1948 and 2020 (Fig. 3e, f). This trend is expected, given the reduction of childhood mortality by advances in health care (28), which reduces the survival benefit of having passed early childhood. Also, parameters that did not follow a monotonous trend, such as parameter *k* (Fig. 3g), showed high consistency across countries. Finally, fitted parameters matched well with commonly used measures of population health: *γ*_*max*_ was highly correlated to childhood mortality under 5 years (Fig. 3h), Parameter *c* was correlated with mortality from chronic disease (Fig. 3i), and *γ*_*max*_ also closely followed life expectancy at birth (Fig. 3j).

In summary, the hDA model describes human mortality by an increasing hazard rate according to the classic GMM, corrected by survival benefits starting either early in life or during adulthood. This extended model fits human mortality data well and offers interpretable parameters.

### Early survival benefits decelerate, and late benefits accelerate the evolution of slower aging in the hDA model

To investigate how the magnitude of early and late survival benefits affected the evolution of aging, we ran agent-based simulations of the hDA model with initial parameters obtained from fits to human mortality data. We used the same simulation design as for the minimal DA model (Fig. 1), except that sexual reproduction was age-dependent, following a Brass polynomial, matching human reproduction dynamics (30).

Similar to the minimal DA model, the aging rate rapidly declined at the start of the simulations in the extended hDA model (Fig. 4), followed by deceleration and an eventual plateau. Increasing *γ*_*max*_ or *λ*_*max*_ elevated this plateau. This behavior is expected, given that the reproduction span in the hDA model is finite. Therefore, reduced extrinsic mortality elevates the threshold of the aging rate below which mortality during the reproductive period reaches near-zero, diminishing the benefit of further slow-down of aging.

**Figure 4.**
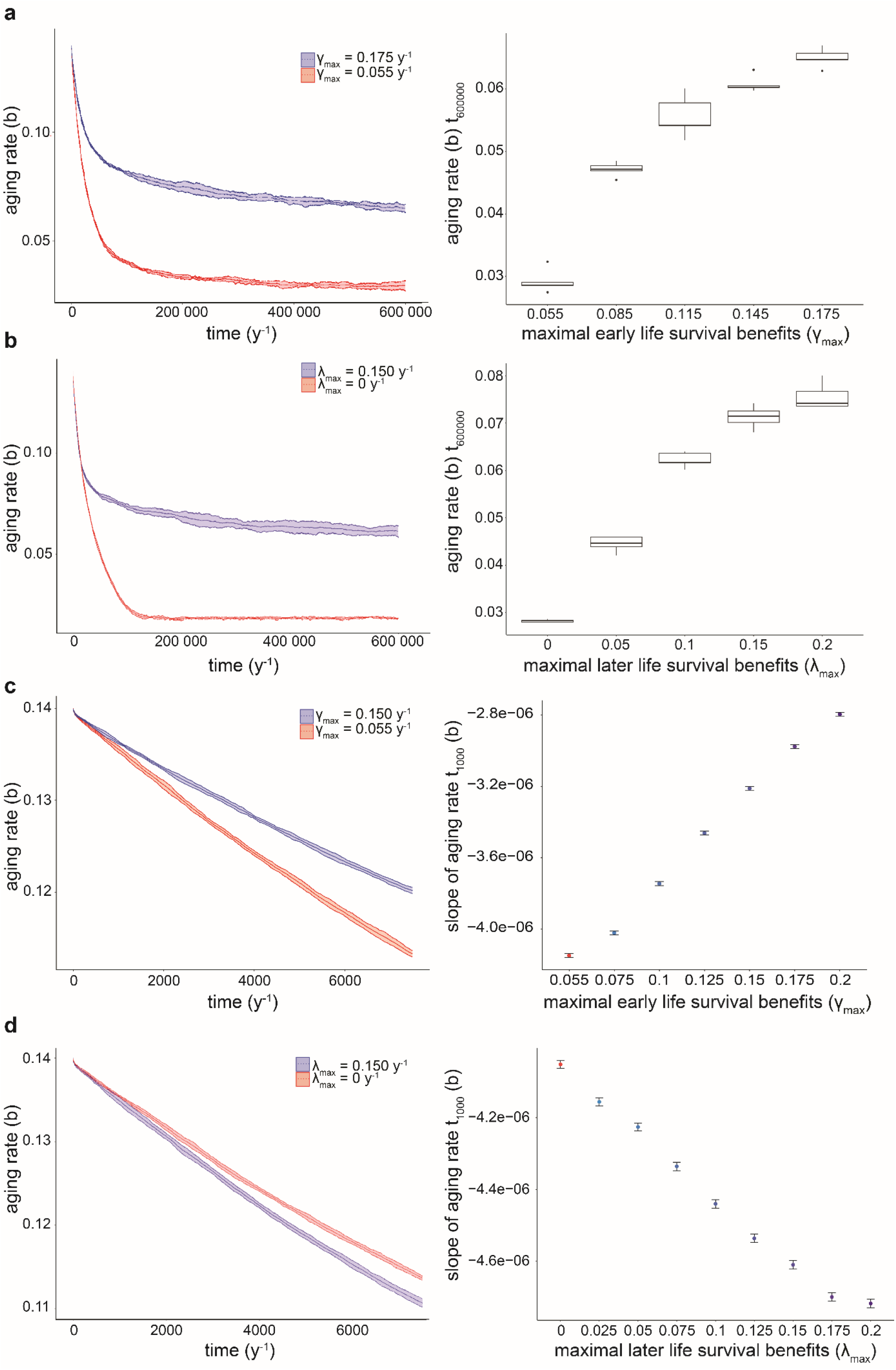
Early survival benefits decelerate, and late survival benefits accelerate the evolution of slower aging in the hDA model. **a**. left: Evolution of aging rate in the simulation of sexually reproducing individuals for two values of *γ*_*max*_. Starting aging rate *b*_*0*_ = 0.15 y^−1^. Fertility function used: Brass polynomial, based on ref. (30). Other life history parameters: fits of the hDA model to mortality data from United Kingdom 1953. right: mean difference in aging rate after 600’000 timesteps (n = 5 independent simulations, with m = 10’000 individuals) as a function of *γ*_*max*_. **b**. as a, but for *λ*_*max*_. **c**. left: as **a** but shorter timescale. right: mean difference in aging rate after 1’000 timesteps (n = 50 independent simulations, with m = 10 000 individuals). **d**. as **c**, but for *λ*_*max*_. Conclusions were not affected by assortative mating, different reproduction functions for males and females, and for asexual simulations (Supplemental Fig. 3). Simulations with *γ*_*max*_ < 0.055 y^−1^ lead to population extinction and are not shown in **a** and **c**.

However, unlike the plateau, the rate of initial decline in the aging rate was differently impacted by *γ*_*max*_ than by *λ*_*max*_. Higher *γ*_*max*_, representing early survival benefits, decelerated the evolution of slower aging, whereas higher *λ*_*max*_, representing late survival benefits, accelerated this evolution (Fig. 4c, d). This behavior of the hDA model is consistent with that of the minimal DA model (Fig. 1). Equivalent results were also obtained in simulations using asexual and age-independent, constant, reproduction, except that the effect on the plateau was less pronounced (Supplemental Fig. 3). This consistency suggests qualitative robustness of our conclusions to using other fertility functions or asexual reproduction.

In summary, simulations of the hDA model confirm a crucial impact of the timing of mortality during development on the evolution of aging that is consistent with conclusions from the minimal DA model.

### Low mortality during reproduction weakens the selection of slower aging

To validate results from agent-based simulations, we computed the intrinsic rate of population increase *r* by the Euler-Lotka equation (24) over a wide range of *λ*_*max*_ and *γ*_*max*_. These calculations yielded a fitness landscape that plateaued at low aging rate *b*, and at high *λ*_*max*_ or *γ*_*max*_ (Fig. 5a, b) for all fertility function tested (Supplemental Figs. 4 and 5). Consistent with agent-based simulations, the slowest rate of aging that could evolve, marked by the edge of the plateau, increased for larger survival benefits *λ*_*max*_ and *γ*_*max*_ (Fig. 5a, b; Supplemental Figs. 4 and 5).

**Figure 5.**
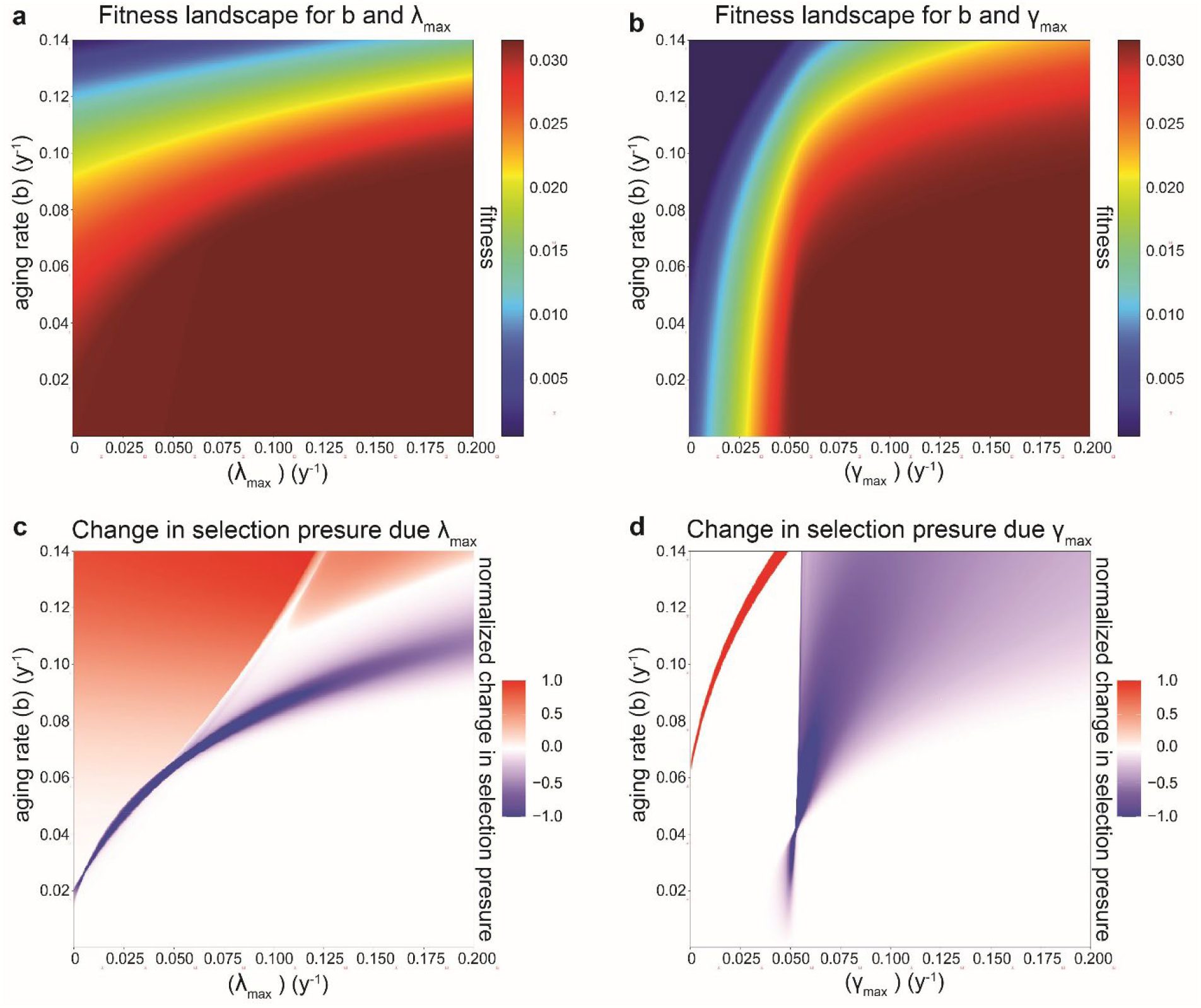
Fitness landscape distinguishes regions where developmental processes accelerate or constrain the evolution of slower aging. **a**. fitness landscape for combination of different aging rates (*b*) and late developmental processes (*λ*_max_). **b**. same as a, but for *γ*_*max*_. **c**. numerical approximation of derivative of selection pressure for slower aging (Δr/Δb) with respect to *λ*_*max*_. The red and blue color scale indicates normalization to the maximal positive or negative value, respectively. **d**. same as **c**, but for *γ*_*max*_

From this landscape, we estimated the selection strength for slower aging, as the change in fitness σ = Δr/Δb resulting from a 5% reduction in the rate of aging *b* (Supplemental Fig. 6) and computed how this selection pressure changed with *λ*_*max*_ and *γ*_*max*_ by computing Δσ/Δ*λ*_max_ and Δσ/Δ*γ*_max_ (Fig. 5c, d, Supplemental Figs. 4 and 5). Across a wide range of parameters outside of the fitness plateau, increasing *λ*_*max*_ increased the selection pressure for slower aging (red area in Fig. 5c). Conversely, increasing *γ*_*max*_ decreased the selection pressure for most values outside of the fitness plateau (blue area in Fig. 5d). The only parameter regions where *λ*_*max*_ and *γ*_*max*_ did not affect *σ*, or had opposite effects, were those with very low total lifetime reproduction or very low mortality during the reproductive period.

In summary, deterministic calculations confirm agent-based simulations for the biologically relevant parameter space, showing that the survival benefits of early and late developmental processes have opposing effects on the evolution of aging over a wide range of parameters outside of the fitness plateaus. Early survival benefits decelerate the evolution of slower aging, whereas late-acting benefits accelerate it.

## Discussion

While early verbal arguments by Williams suggested that extrinsic mortality leads to the evolution of faster aging (10), subsequent mathematical analysis revealed a more nuanced relationship that is influenced by population density (9,11) and age-dependence (8,14,17). The role of density-dependence has been extensively studied (9,11). However, while age-dependent extrinsic mortality has been acknowledged as being essential for influencing the evolution of aging in the absence of density effects (8), the specific impact of timing has not been systematically investigated. Here, we revealed a consistent pattern: reduced mortality early in life hinders the evolution of slow aging, and reduced mortality later in life accelerates this evolution.

The impact of survival benefits on the evolution of aging can be understood by the separating two distinct effects (9): The first effect, following Williams’ reasoning, is that improved survival of extrinsic threats increases the fraction of individuals that live long enough to be affected by aging. As illustrated in Fig. 6a-b, this effect amplifies the relative change in mortality and consequently the lifespan gains from a given reduction in the aging rate, thereby increasing the selective advantage of a slower aging rate. The second, opposing effect is that improved survival accelerates population growth, which in turn diminishes the relative fitness value of individuals reaching an old age (11). This effect reduces the selective benefit of slower aging.

**Figure 6.**
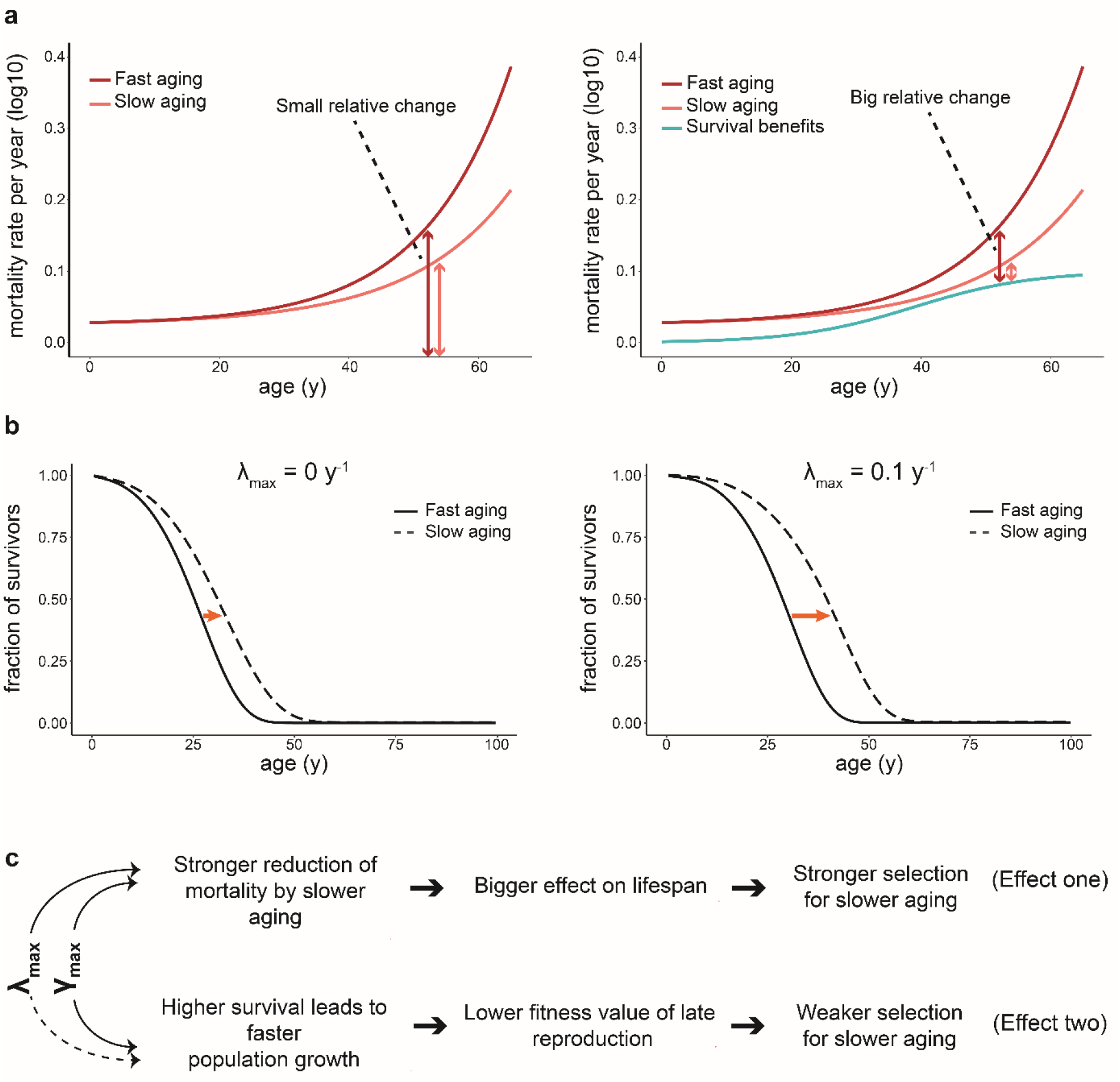
Differential impact on population growth and lifespan gain explains opposite effect of early and late survival benefits for slow aging. **a**. Example of mortality rates for the GMM for two different aging rates (dark and light red), and for developmental survival benefits (cyan, right panel only) as a function of age. Net mortality is the difference between GMM and survival benefits, shown by vertical arrows. Relative change in mortality upon reduced aging rate is larger in the presence of survival benefits (right). **b**. Change in lifespan after reduction of aging rate for scenario with (right) and without (left) survival benefits. fast aging: *b* = 0.085 y^−1^, slow aging: *b* = 0.065 y^−1^. **c**. Summary of differential impact of early and late survival benefits on aging evolution.

When extrinsic mortality is age-independent and in the absence of density effects, these two evolutionary effects precisely cancel each other out, resulting in no net selection on aging (9,11,14). However, when extrinsic mortality is modulated with age, one of the two effects can dominate, generating a net selection pressure (11,20,21,31). Both early and late survival benefits increase the fraction of individuals surviving to an old age, which promotes the evolution of slow aging via the first effect. However, because late survival benefits influence only a fraction of the reproductive window, they have a weaker effect on population growth, and thus a smaller contribution to the second, counteracting effect. Consequently, late survival benefits lead to a net increase in the selection for slower aging, while early benefits tip the balance towards weaker selection for slower aging (Fig. 6c).

The hDA model does not specify biological mechanisms responsible for the survival benefits and instead uses phenomenological functions to capture age-dependent mortality based on empirical human data. Several known age-dependent processes could underlie such survival benefits. Early benefits are likely driven by postnatal growth and development, while late survival benefits may stem from learning, adaptive immunity, or, in some species, complex social behaviors. The wide variety of such processes across taxa highlights the importance of considering developmentally controlled survival benefits to understand the diversity of aging rates.

Our findings on the timing of extrinsic mortality offer a new perspective on experimental evolution studies that have yielded unexpected results. For example, a landmark study in guppies found that populations under high predation pressure evolved slower aging, rather than faster aging (32), contrasting predictions from classic theory (10). Previous explanations invoked density dependence (12,33), environmental and genetic variation (34), or changes in food abundance (35). Our model provides an alternative interpretation: because the predators used in the study predominantly affected young individuals (36), the increase in mortality was likely concentrated early in life. According to our model, such early-life mortality reduces the selection for slow aging, aligning with the experimental observations. This matching between experiment and theory underlines the applicable significance of our model.

In addition to distinguishing the effects on the speed of aging evolution (i.e., the slope of the aging rate), our model also makes predictions on the slowest rate of aging that could evolve in the absence of tradeoffs (i.e., the aging plateau). Our interpretation for the emergence of the plateau is that when survival benefits reduce mortality so strongly that aging has no impact on the reproductive output, then the selection pressure for slower aging diminishes. Consequently, our theory predicts that evolution of negligible senescence can only occur when developmental survival benefits are very low, offering an explanation why negligible senescence is so rare in nature.

## Conclusions

Our study shows that the timing of changes in extrinsic mortality plays a crucial role in shaping the evolutionary pressures on aging. Specifically, reducing mortality early in life diminishes selection for slower aging, whereas reducing mortality later in life strengthens it. This time-dependence arises from the interplay between two opposing forces: on one hand, lower mortality increases the number of individuals reaching old age, thereby intensifying selection for longevity; on the other hand, it also accelerates population growth, which weakens this selection pressure. Early-life mortality reductions have a stronger impact on the population growth rate and thus reduce the evolutionary advantage of slower aging. In contrast, late-life mortality reductions primarily increase the proportion of long-lived individuals, enhancing selection for slow aging. Recognizing this nuanced effect of mortality timing helps explain previously puzzling outcomes in studies of extrinsic mortality and aging evolution.

## Methods

### Minimal DA model

Mortality increases exponentially with time according to the Gompertz law

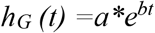

Survival benefits *β* were a step function that decreases mortality by 20% from age *d* onwards, yielding an overall mortality

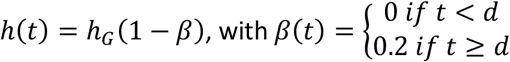

Reproduction was sexual with constant fertility of one offspring per time step between ages 25 to 100 and zero fertility outside of this interval.

### Details of the hDA model

The hDA model describes the hazard rate *h(t)* as the sum of effects of aging (α), early life mortality benefits (*γ*) and later life mortality benefits (*λ*) on survival, each of which has a distinct time dependence:

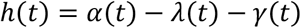

aging follows the Gompertz-Makeham function (23), a commonly used model of aging:

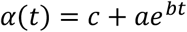

*c, a*, and *b* are constant parameters, and *t* represents time. The parameter *b* is called the aging rate, the exponential rate at which mortality due to aging increases with time (i.e., in the absence of developmental survival benefits).

Later life mortality benefits reduce mortality with age by a logistic function which plateaus to a maximal benefit *λ*_*max*_:

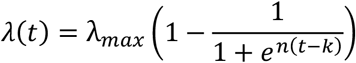

*k* is the age at which the later life benefits reach half their maximum. *n* determines the steepness of the curve.

We modeled the early life benefits by a decelerating function.

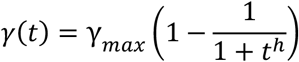

*γ*_max_ corresponds to the maximal early survival benefits and *h* relates to the steepness of the curve.

### Mortality data

Data on human mortality rates were acquired from the human mortality database (37) on 20 July 2023. We have analyzed the population data (cross-sectional). Analysis was restricted to mortality data after 1948. We focused on the 8 countries with a population of over 10 million in 1948. Population data was taken from 1948 to 2020 for the same countries.

### Fitting the hDA model

The Least-Squares method (implemented in the scipy.optimize Python package) was used to fit the parameters of the realistic DA model to log-transformed mortality data.

### Computation of fitness functions and the benefit of slower aging

We used the intrinsic rate of population increase *r* as a measure of Darwinian fitness, defined by the Euler-Lotka equation (24)

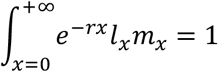

*l*_*x*_ is the likelihood of survival to age *x. m*_*x*_ corresponds to the number of offspring produced by individuals of age *x*. Based on the hDA model, survivorship *l*_*x*_ was calculated as:

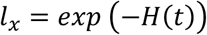

With *H* corresponding to the cumulative hazard rate.

For *m*_*x*_, we used a Brass Polynomial defined as:

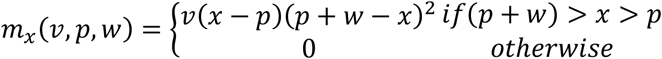

For models shown in Supplemental data, we used constant (*m*_*x*_ = *m*_*0*_), increasing (*m*_*x*_ = *m*_0_ + 0.1 *x*), or decreasing (*m*_*x*_= *m*_0_ − 0.1*x*) fertility functions. For the decreasing function, fertility was not allowed to go below 0.

Fitness landscapes in Fig. 4 were based on parameters from fits to United Kingdom population mortality data from 1953. Fitness landscapes produced using parameters from other countries and years were very similar. Landscapes were generated using Rust.

Numerical integration was performed using the peroxide Rust crate.

### Agent-based simulations

For all agent-based simulations, time was discrete, with each time step corresponding to the age increment of 1 and with all relevant rates and probabilities defined on a per-time-step basis. Each simulation was run in 5 replicates unless stated otherwise in the main text. Simulations were carried out with a population of 10’000 individuals.

In all simulations, individual agents underwent repeated rounds of death, reproduction, and mutation. During the death part of the simulation step, the cumulative change of dying during the current step was calculated for each individual, and a Bernoulli trial determined survival. During the reproduction step, offspring were created, with a 2% chance of a mutation to alter the rate of aging (i.e, parameter *b)*. Out of the created offspring, a random subset of offspring was selected to fill free spaces up to the population limit.

### Details of agent-based simulation of the hDA model

Parameters were taken from fits to United Kingdom population mortality from 1953. Conclusions were robust to using parameters from fits to cohort or population form data from other countries and years.

Each agent was represented by four attributes: its sex, its age *x*, its aging rate *b*, and its maximum survival benefits (*λ*_*max*_ and *γ*_*max*_). When initializing the simulation, each agent was assigned *x* and *b* drawn from a normal distribution (mean(*x*) = 20, SD(*x*) = 10; mean (*b*) = 0.14, SD (*b*) = 0.005). *λ*_*max*_ was initialized as a constant for all individuals. The population size was set to 10’000.

For every round of the simulation, cumulative probability of death was computed based on the hDA model for each agent. Survivorship was determined by a Bernoulli trial, followed by either increasing the age by 1 or removing the agent from the simulation.

Reproduction was modeled by the Brass Polynomial as described above using parameters obtained from human fertility data of men and women (30). To optimize computations, simulations were discontinued for agents whose age was above the fertile period (*w* + *p* from the Brass polynomial). Simulations, where post-reproductive individuals were not removed, led to qualitatively identical conclusions. However, in such a setting, the eventual existence of non-dying but no longer reproducing individuals in the control condition (*λ*_*max*_= 0) led to a sudden drop of the mean *b* to zero (Supplemental Fig. 7). The probability *p* of an agent to reproduce in each round was calculated from the brass polynomial by:

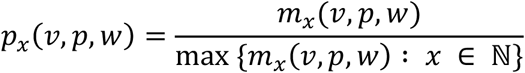

All agents were then randomly paired as male-female couples. If both agents of a couple were able to reproduce, they produced a new agent of random sex. Simulations with constant fertility rate and assortative mating with regard to the age of the agent led to identical conclusions (Supplemental Fig. 3). The aging rate *b* of the new agent was assigned the mean *b* of its parents. 2% of all newly created agents were then randomly selected to mutate and assigned an aging rate *b* drawn from a normal distribution with mean being its pre-mutation value and SD = 0.012.

Parameters used in the simulations are summarized in Supplemental Table 1. Simulations were all run using Rust.

## Declarations

Code used for this study is available at https://github.com/PeterLenart/Learning_accelerates_aging.git

Human mortality data analysed in the study were obtained on 24. May 2022 from The Human Mortality Database (https://www.mortality.org/)

## Authors’ contributions

PL formulated the research problem, created the model, designed and ran the agent-based simulations, executed fitness function computations, and wrote the manuscript. SP, improved the model and algorithms for fitting, computation of fitness landscapes, and agent-based simulations and co-wrote the manuscript. BDT supervised the project and edited the manuscript.

