## supporting_information for "Timing of mortality during development alters the evolution of aging"

**Time-dependent impact of developmental processes on the evolution of aging**

Figure S1.

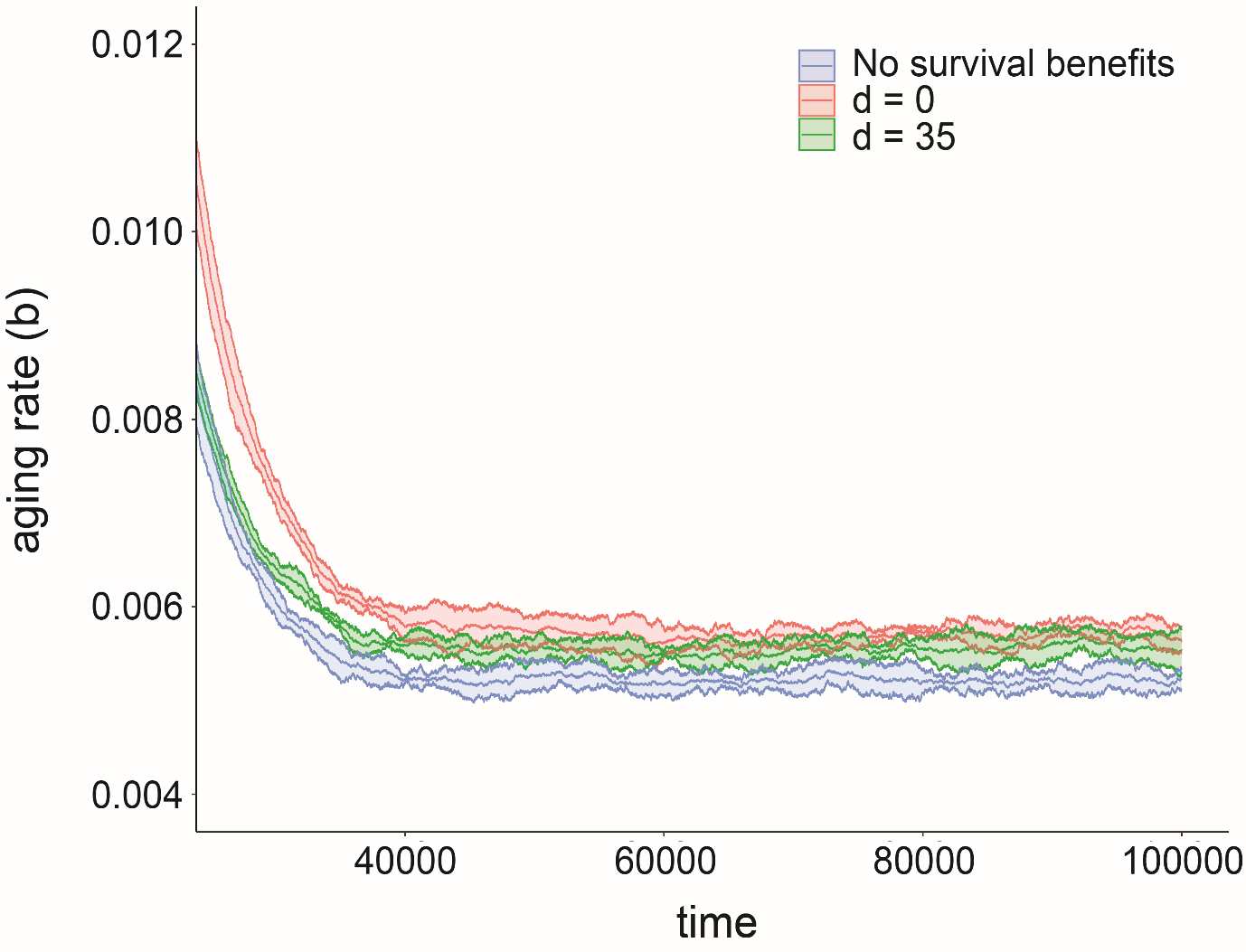

**Figure S1. The evolution of aging rates in the DA model plateaus in the long term.** Same simulation as shown in Figure 1a but for time steps 20’000 to 100’000 to show plateau of aging rate evolution. Colours represent populations with different timings of survival benefits early (red) or late (green) in life, compared to a population without developmentally controlled survival benefits (blue). The middle line of each trajectory shows the mean of 5 replicates (10‘000 agents each). The shaded area represents a 95% confidence interval.

Figure S2.

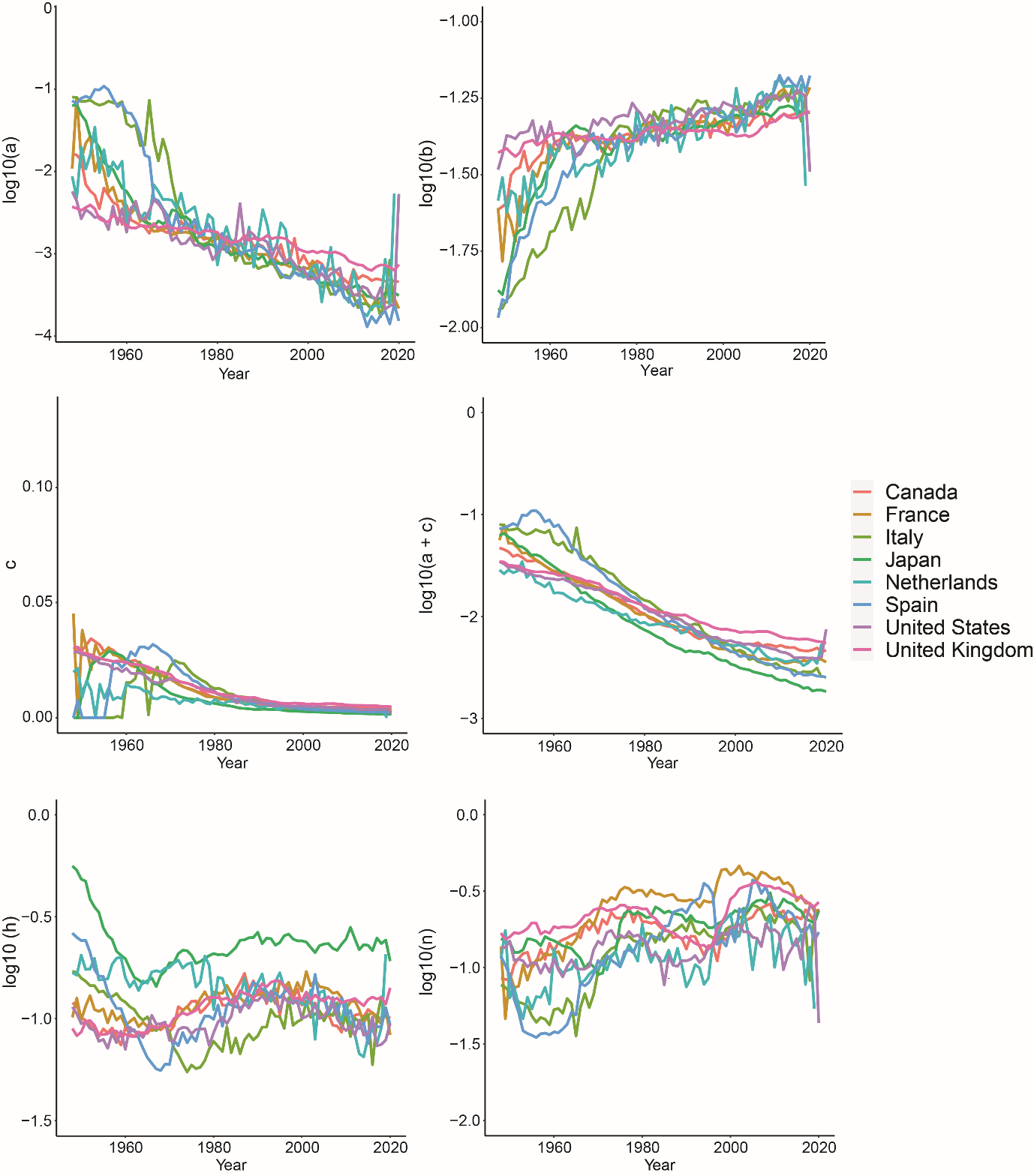

**Figure S2. Parameters of the hDA model follow systematic trends over time**

Panels show the change of parameters of the hDA model obtained from fits to mortality data from different years and countries, as indicated by colors in the legend.

Figure S3.

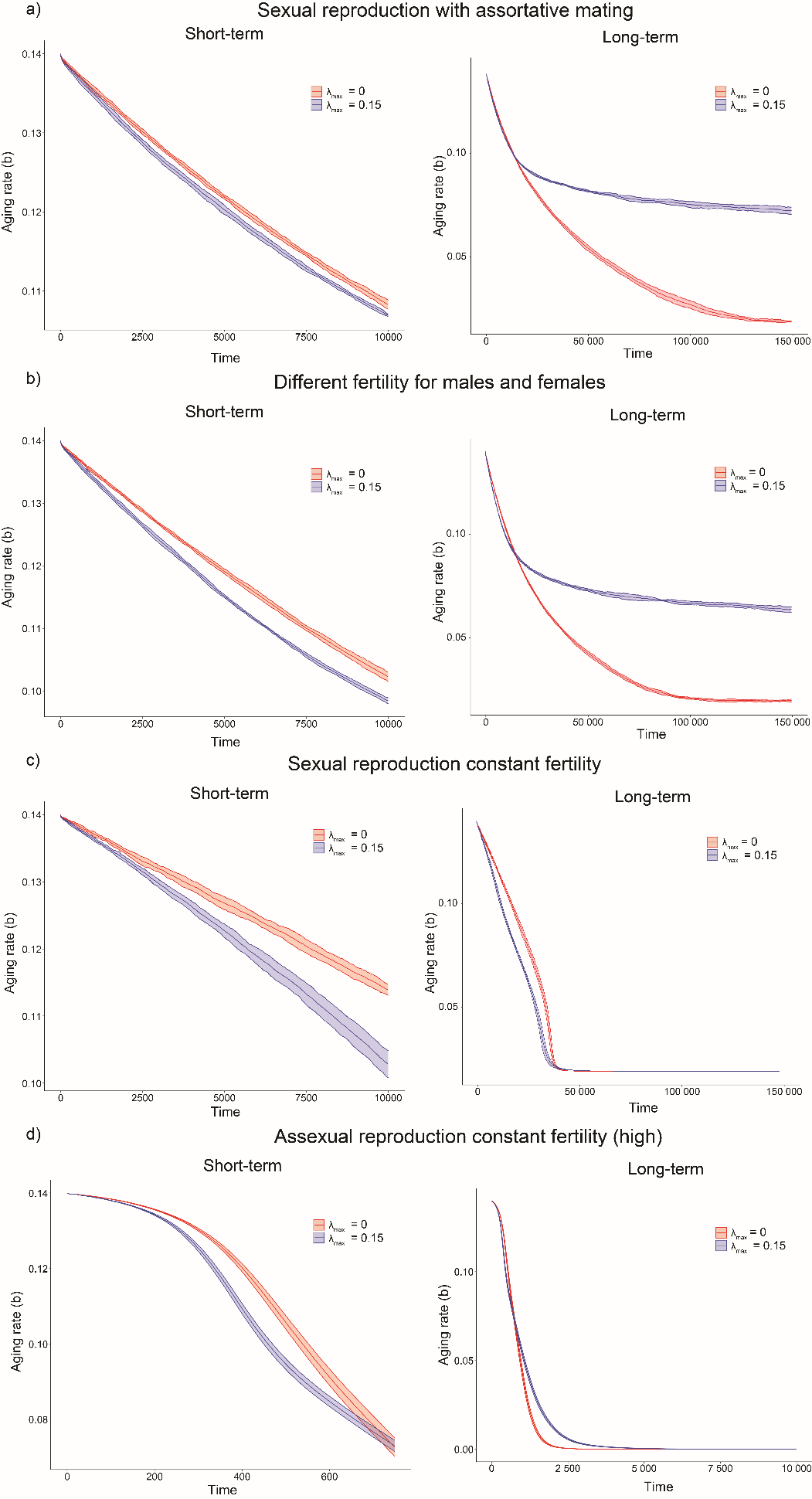

**Figure S3. Impact of developmental survival benefits for populations with different reproductive functions and sexuality.** Agent-based simulations of evolution under the hDA model and asexual reproduction with indicated values for $\lambda_{max}=0.15$(blue) or $\lambda_{max}$ = 0 (red). **a.** same as sexual simulation shown in Fig. 4, but agents mate only with individuals of the same age **b.** male fertility declines slower than female (reproductive period from 14.8 to 47.8 compared to a reproductive period of 14.8 to 32.8 in females) see Table S1 **c.** sexual reproduction without fertility decline with age **d.** asexual simulation with constant fertility setting. Detailed simulations parameters are described in methods. Middle line: mean of 20 simulations (10‘000 individuals each) Filled area: 95% confidence interval. For all simulations, the initial decline in the aging rate is faster in the presence of survival benefits (blue line below red line). Over time, evolution slows down more strongly in the presence of survival benefits (blue line is above red line),

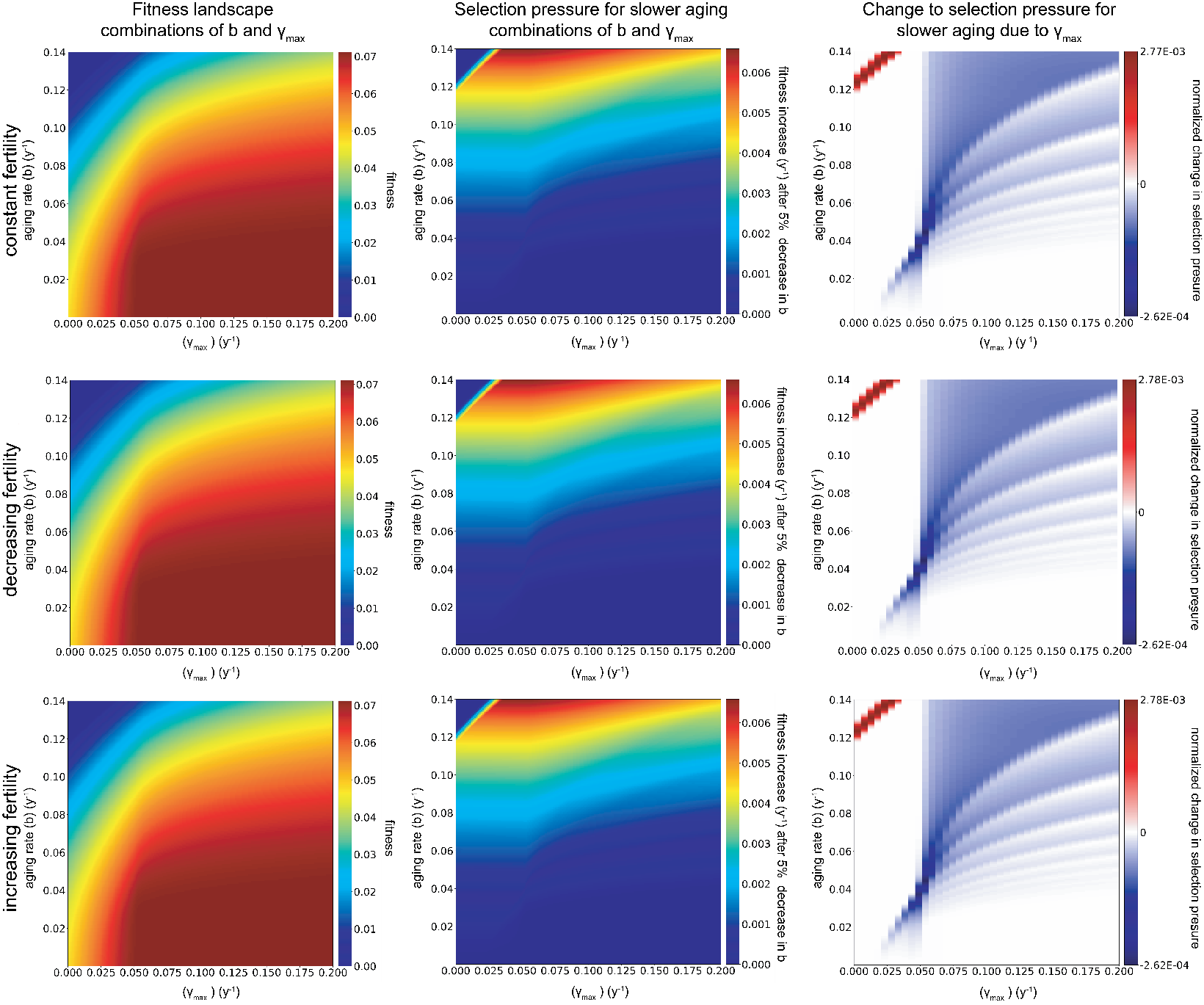
Figure S4.

**Figure S4. Early developmental survival benefits increase the fitness benefit of slower aging for decreasing, increasing, and constant fertility functions.** Left: fitness (intrinsic growth rate *r*) as a function of aging rate *b* and late survival benefit $\gamma_{max}$. Middle: selective pressure for slower aging measured as fitness benefit of reducing the aging rate by 5% as a function of maximal early survival benefit $\gamma_{max}$ and initial aging rate *b_0_*. Right: Numerical approximation of the derivative of selection pressure change with respect to *γ_max_* . Blue indicates that increasing survival benefits accelerate the evolution of slower aging. Red indicates that survival benefits reduce the selection pressure. Negative and positive selection pressures were normalized to their respective maximum for display. First row, constant fertility function ($\boldsymbol{m}_{\boldsymbol{x}}=\boldsymbol{c}$, with c = 0.01). Second row, fertility function decreases linearly with age ($\boldsymbol{m}_{\boldsymbol{x}}\boldsymbol{= ax +b}$, with a = -0.001 and b = 0.01). In the third row, the fertility function increases with age $\boldsymbol{(m}_{\boldsymbol{x}}\boldsymbol{= ax+b,}$ with a = 0.001 and b = 0.01). $\boldsymbol{m}_{\boldsymbol{x}}$ was set to 0 for $\boldsymbol{x<25.}$

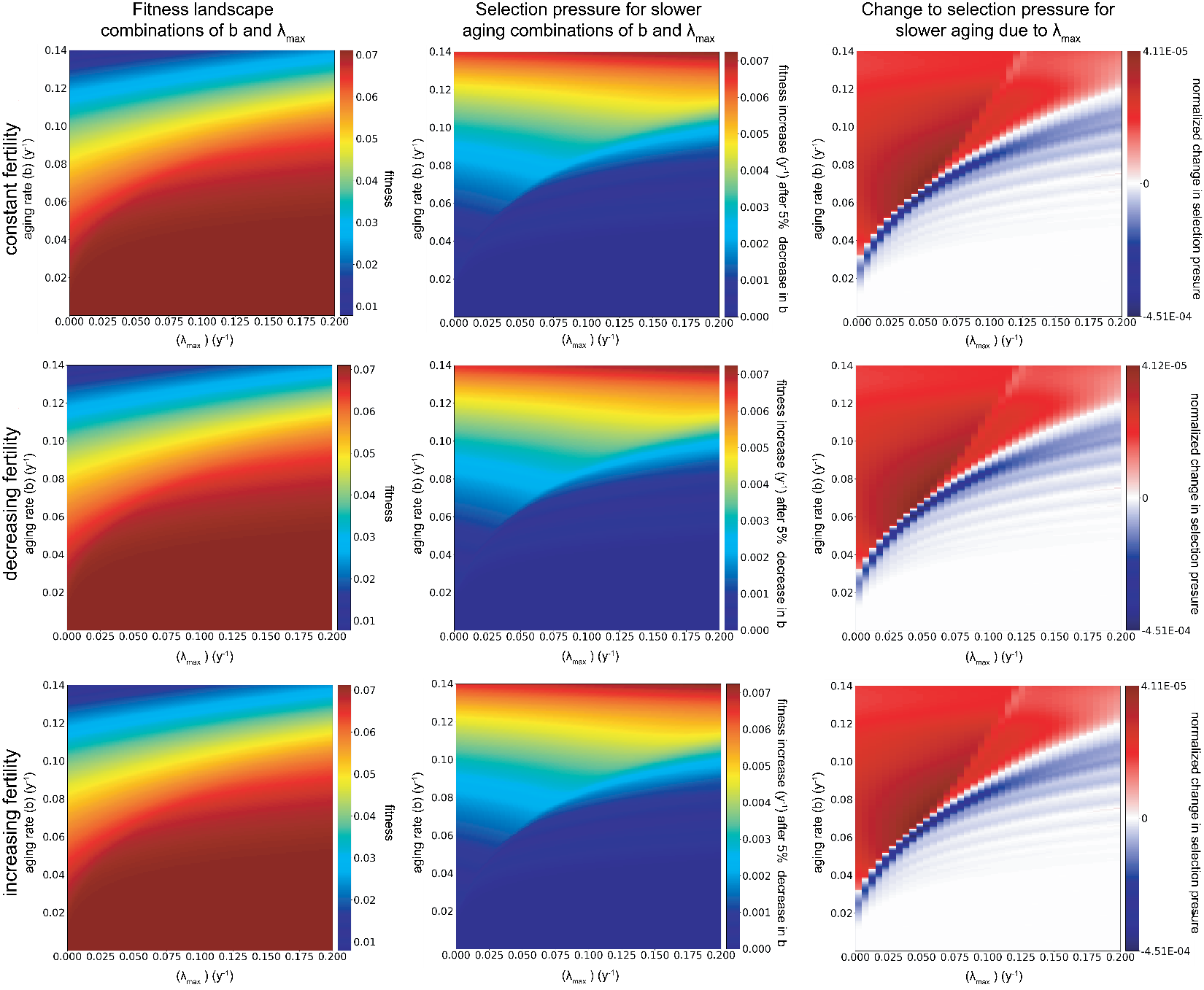
Figure S5.

**Figure S5. Late developmental survival benefits increase the fitness benefit of slower aging for decreasing, increasing, and constant fertility functions.** Left: fitness (intrinsic growth rate *r*) as a function of aging rate *b* and late survival benefit $\lambda_{max}$. Middle: selective pressure for slower aging measured as the fitness benefit of reducing the aging rate by 5% as a function of maximal early survival benefit $\lambda_{max}$ and initial aging rate *b_0_*. Right: Numerical approximation of the derivative of selection pressure change with respect to $\lambda_{max}$. Blue indicates that increasing survival benefits accelerate the evolution of slower aging. Red indicates that survival benefits reduce the selection pressure. Negative and positive selection pressures were normalized to their respective maximum for the display. First row, constant fertility function ($\boldsymbol{m}_{\boldsymbol{x}}=\boldsymbol{c}$, with c = 0.01). Second row, fertility function decreases linearly with age ($\boldsymbol{m}_{\boldsymbol{x}}\boldsymbol{= ax +b}$, with a = -0.001 and b = 0.01). In the third row, the fertility function increases with age $\boldsymbol{(m}_{\boldsymbol{x}}\boldsymbol{= ax+b,}$ with a = 0.001 and b = 0.01). $\boldsymbol{m}_{\boldsymbol{x}}$ was set to 0 for $\boldsymbol{x<25.}$

Figure S6.

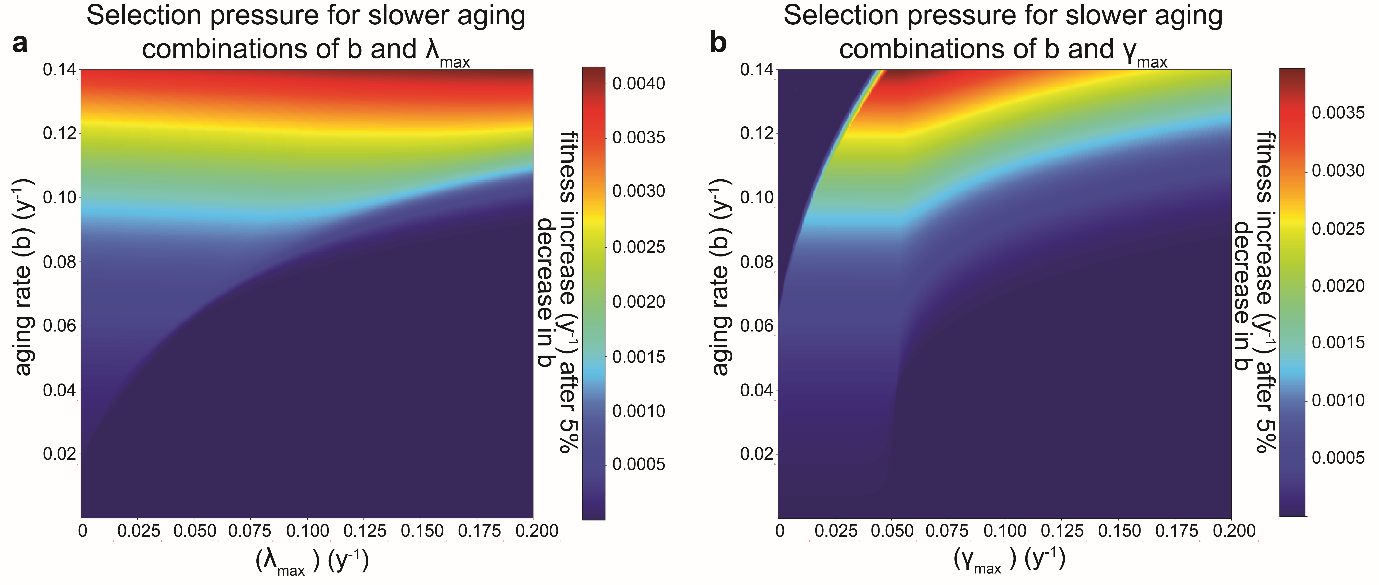

**Figure S6. Selection pressure for slower aging changes with survival benefits**. **a**. selective pressure for slower aging as a function of $\lambda_{max}$ and the initial rate of aging *b_0_*. **b**. as a, but for γ_max_. Selective pressure was calculated as the absolute fitness benefit in intrinsic rate of population increase *r* from a 5% reduction in the rate of aging.

Figure S7.

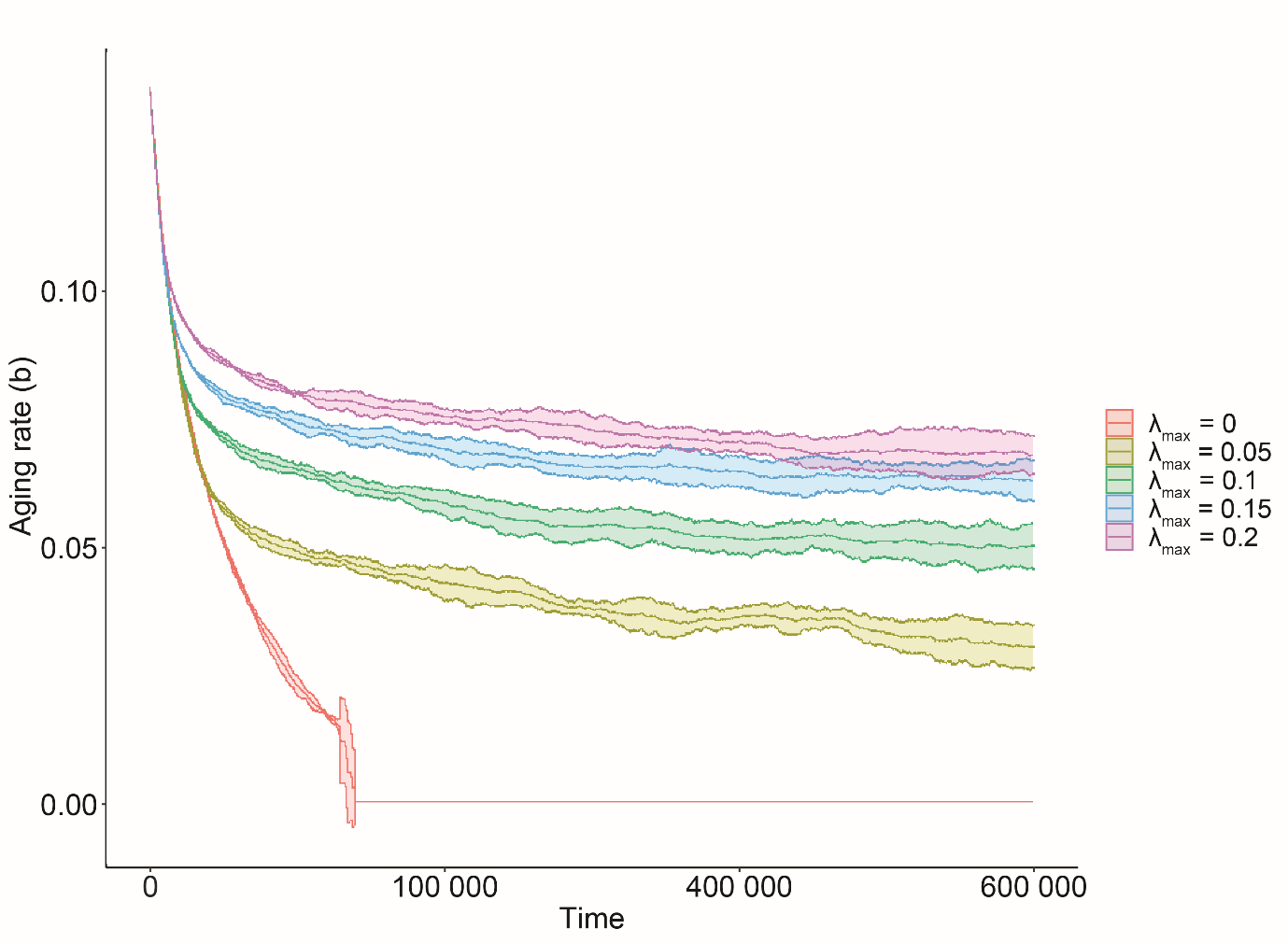

**Figure S7. Simulation behavior is robust to omitting the removal of the post-reproduction individuals and growth in population size.** Agent**-**based simulations were conducted using sexual reproduction as described in Figure 3, except that post-reproductive individuals were not removed from the simulations. The middle line of each trajectory shows the mean of 5 replicates (10‘000 agents each). The shaded area represents a 95% confidence interval.
